# Atlas of Fetal Metabolism During Mid-To-Late Gestation and Diabetic Pregnancy

**DOI:** 10.1101/2023.03.16.532852

**Authors:** Cesar A Perez-Ramirez, Haruko Nakano, Richard C Law, Nedas Matulionis, Jennifer Thompson, Andrew Pfeiffer, Junyoung O Park, Atsushi Nakano, Heather R Christofk

## Abstract

Mounting evidence supports an instructive role for metabolism in stem cell fate decisions. However, much is yet unknown about how fetal metabolism changes during mammalian development and how altered maternal metabolism shapes fetal metabolism. Here, we present a descriptive atlas of *in vivo* fetal murine metabolism during mid-to-late gestation in normal and diabetic pregnancy. Using ^13^C-glucose and LC-MS, we profiled the metabolism of fetal brains, hearts, livers, and placentas harvested from pregnant dams between embryonic days (E)10.5 and 18.5. Comparative analysis of our large metabolomics dataset revealed metabolic features specific to fetal tissues developed under a hyperglycemic environment as well as metabolic signatures that may denote developmental transitions during euglycemic development. We observed sorbitol accumulation in fetal tissues and altered neurotransmitter levels in fetal brains isolated from dams with maternal hyperglycemia. Tracing ^13^C-glucose revealed disparate nutrient sourcing in fetuses depending on maternal glycemic states. Regardless of glycemic state, histidine-derived metabolites accumulated during late development in fetal tissues and maternal plasma. Our rich dataset presents a comprehensive overview of *in vivo* fetal tissue metabolism and alterations occurring as a result of maternal hyperglycemia.

## INTRODUCTION

Maternal nutrition and fetal development are inseparable. Dietary interventions during pregnancy promote maternal health and reduce the incidence of fetal developmental defects. For instance, iron supplementation prevents maternal anemia during pregnancy (Georgieff, 2020), and folic acid supplementation reduces the incidence of fetal neural tube defects (Scholl and Johnson, 2000). However, not much is known about how fetal metabolism is shaped during development, nor how prevalent clinical conditions affecting maternal metabolism impact developing fetuses. Recent studies have begun to characterize the metabolic plasticity and compartmentalization in the mouse pre-implantation embryo (Sharpley et al., 2021) and during embryonic day (E)9.5-13.5 (Solmonson et al., 2022). But fetal metabolism during mid-to-late gestation, when the fetus nears completion of its independent circulatory system, remains unexplored.

Maternal diabetes is a growing clinical problem in the US and worldwide: the rate of diabetes cases amongst pregnant individuals per 1,000 live births in the US rose from 47.6 in 2011 to 63.5 in 2019 (Shah et al., 2021). This adverse trend is due to an increased incidence of gestational diabetes and type 2 diabetes in younger patients prior to pregnancy. Maternal hyperglycemia is associated with a four-fold increased risk of congenital heart defects and increased risk of neurodevelopmental defects (Ornoy et al., 2021; Øyen et al., 2016; Tinker et al., 2020). Previously, we found that high glucose levels impair cardiac maturation *in vitro* through excessive nucleotide metabolism (Nakano et al., 2017). However, the *in vivo* impact of maternal hyperglycemia on the metabolism of the developing fetus remains unknown and may shed light mechanistically on the cause of congenital defects in diabetic pregnancy.

Remarkable studies in mice have provided an increasingly detailed view of mammalian development from the perspective of genomics and transcriptomics (Chen et al., 2022; Gorkin et al., 2020; He et al., 2020). Here we provide an additional layer to the understanding of mammalian development by quantifying the metabolic landscape of the developing fetus at a multi-tissue level and examining how maternal hyperglycemia shapes this landscape. Using LC-MS and [U-^13^C]glucose, we measured the metabolic profile of fetal tissues (placenta, heart, liver, brain) dissected from pregnant mice during mid-to-late gestation. We present an expansive metabolomic atlas of fetal tissues that offers insights into metabolite level dynamics and pathway utilization under euglycemia and hyperglycemia. Our rich dataset reveals metabolic signatures that reflect the impact high glucose *in utero* has on fetal tissue metabolism and can be leveraged toward the design of improved strategies to meet the nutritional needs of the fetus.

## RESULTS

### Fetal metabolomics in euglycemia and hyperglycemia

To model diabetic pregnancy, we used the hyperglycemic Akita mouse, which produces mildly defective but surviving offspring through the mid- and late-gestation periods. These mice harbor a heterozygous mutation in the Ins2 gene, leading to the development of adult-onset diabetes (Yoshioka et al., 1997). In the C57BL/6 background, Akita dams were crossed with wildtype males to create a diabetic pregnancy condition in which fetuses are exposed to a hyperglycemic environment (Nakano et al., 2017). Offspring of Akita dams showed some heart and neural tube defects; however, most defects are mild with the majority being suitable for LC-MS-based metabolomic analysis (Figure 1A and S1A).

**Figure 1.**
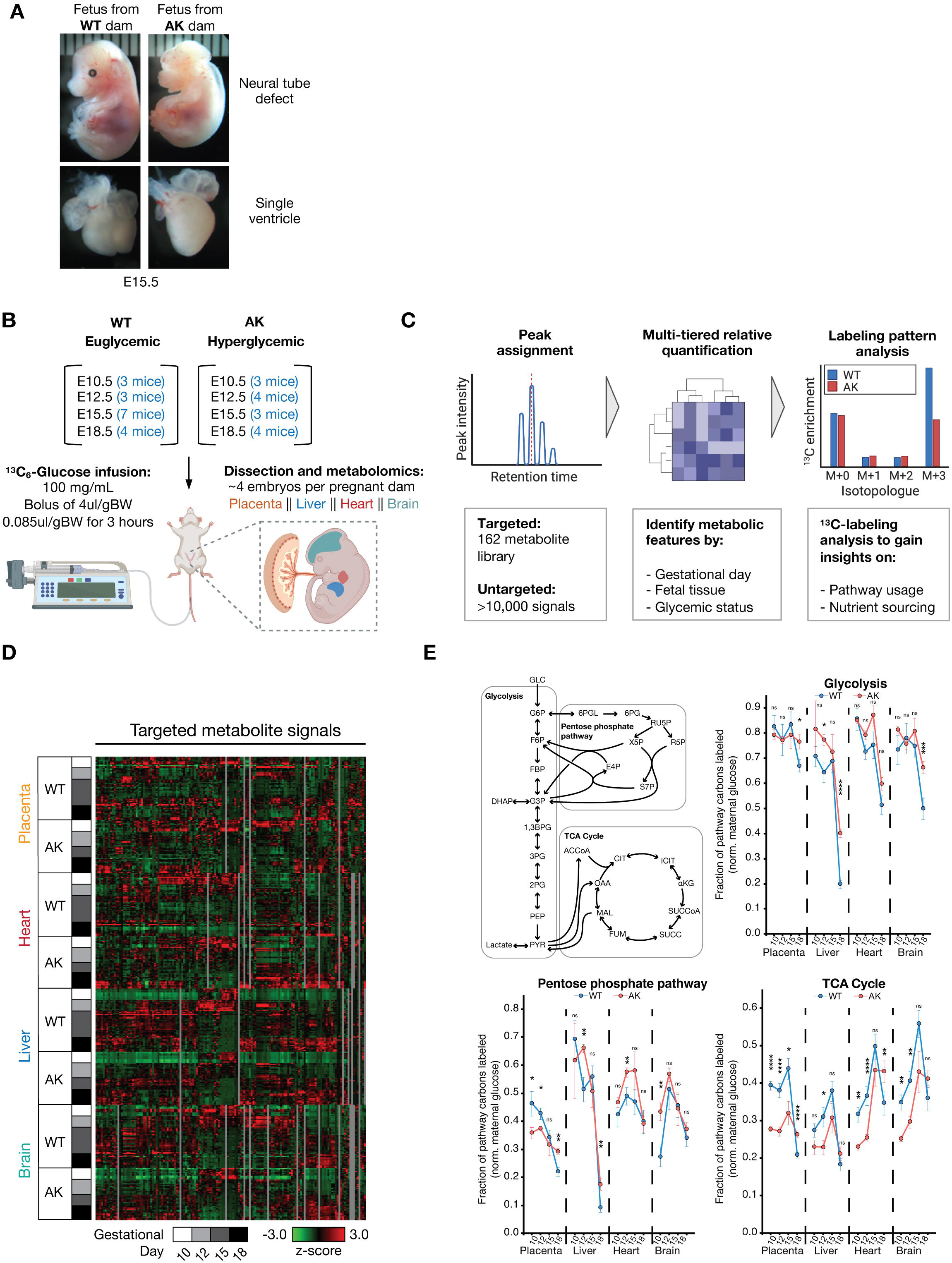
Metabolomic platform to assess fetal metabolism during mid-to-late gestation and diabetic pregnancy. (A) Images of fetuses harvested from wildtype (WT) and Akita (AK) dams at embryonic day 15.5 highlighting neural tube defect and single ventricle defect in fetuses from Akita dams. (B) Schematic of experimental setup involving timed pregnancies in WT and AK dams, [U-^13^C_6_] glucose infusion parameters, and tissues collected. (C) Schematic of LC-MS-based metabolomic data analysis approach. (D) Heatmap representation of targeted metabolite analysis across placenta, heart, liver, and brain in fetal tissues isolated from WT and AK dams across multiple timepoints during development. Plotted data represents z-score. (E) Fraction of pathway carbons labeled normalized to the level of maternal glucose labeling. Fractional labeling of carbons from metabolites in a pathway (glycolysis, pentose phosphate pathway, TCA cycle) was averaged weighted by number of carbons per metabolite to indicate whole pathway labeling at different timepoints during mid-to-late gestation. Statistical analyses were determined using two-tailed *t*-tests.

We set out to map the metabolomic profile of fetal tissues collected from pregnant Akita dams versus wildtype dams throughout development, specifically during mid-to-late gestation. To accomplish this goal, timed mouse pregnancies were set to stages E10.5, E12.5, E15.5, and E18.5. Both wildtype and diabetic (Akita) dams were fasted overnight and infused with [U-^13^C]glucose prior to fetal tissue collection to trace ^13^C through metabolic pathways. Following three hours of tracer infusion, metabolites were extracted from dissected fetal placenta, brain, heart, and liver (Figure 1B). Metabolites from maternal plasma samples were also extracted and analyzed by LC-MS to ensure elevated glucose levels in the Akita dams and similar ^13^C-glucose enrichment in wildtype and Akita dams (Figure S1B and S1C). This experimental setup yielded a high-quality dataset providing three analytical dimensions within the context of development: 1) developmental stage, 2) fetal tissue origin, and 3) maternal glycemic state. Our data analysis pipeline involved both targeted and untargeted metabolomics as well as isotope tracing from [U-^13^C]glucose to provide insights into pathway utilization and nutrient sourcing (Figure 1C). Targeted analysis of metabolomics data involved the cross-referencing to a metabolite library composed of 162 metabolites (Table S1). Across all tissues, gestational stages, and maternal glycemic state, we generated a comprehensive quantification of metabolite levels for further analysis (Figure 1D). Furthermore, we measured the isotopologues of each metabolite to gain further insights into metabolic fluxes (Table S2). Fractional ^13^C enrichment analysis showed disparate activities across developmental stages and metabolic pathways (Figure 1E). In all fetal tissues, glycolytic metabolites were mostly labeled between E10.5 and E15.5, but glycolytic metabolite labeling sharply decreased in E18.5 fetal tissues. Glycolytic metabolites in the liver and the brain of E18.5 fetuses from wildtype dams were significantly less labeled than those from Akita dams, indicating a growing disparity in glycolytic turnover rates depending on maternal glycemic state. The pentose phosphate pathway (PPP) ^13^C enrichment suggested peak activities at different development stages for different tissues: the PPP activity in the placenta and the liver peaked at E10.5 while in the heart and the brain it peaked at E12.5 and E15.5. Multiple nutrients contribute to the TCA cycle metabolite pool. Thus, its ^13^C enrichment indicated the direct and indirect contribution of glucose to the TCA cycle. The fetal tissues from wildtype dams were significantly more labeled in the early E10.5 and E12.5 stages but the labeling in fetal tissues from Akita dams overtook the wildtype counterparts by E18.5, indicative of sustained glucose catabolism in hyperglycemic development.

### Maternal hyperglycemia causes sorbitol accumulation in fetal tissues

Targeted metabolomic analysis of our dataset revealed markedly increased sorbitol levels in fetal tissues harvested from Akita versus wildtype dams (Figure 2A). Sorbitol is generated from glucose through the action of aldose reductase as part of the polyol pathway (Figure 2B). To assess the extent of sorbitol accumulation in fetal tissues harvested from Akita versus wildtype dams, we determined the relative increase in sorbitol levels relative to E10.5 fetal tissues from wildtype dams. We observed increased sorbitol levels in fetal placentas, hearts, livers, and brains exposed to maternal hyperglycemia. Fetal brains isolated from Akita dams during E15.5 and E18.5, in particular, showed the largest sorbitol accumulation relative to E10.5 fetal brains isolated from wildtype dams (Figure 2C). Furthermore, increased sorbitol levels were observed in maternal plasma from E10.5 and E12.5 Akita dams (Figure S2). Sorbitol accumulation in adult diabetic patients can lead to tissue osmotic stress and contributes to the damage that occurs in insulin-independent tissues such as retina, kidney, and nerves (Brownlee, 2001). Our data revealed that fetuses exposed to maternal hyperglycemia are not protected from sorbitol accumulation and suggested that sorbitol accumulation during fetal development may contribute to the higher incidence of developmental defects observed during diabetic pregnancy.

**Figure 2.**
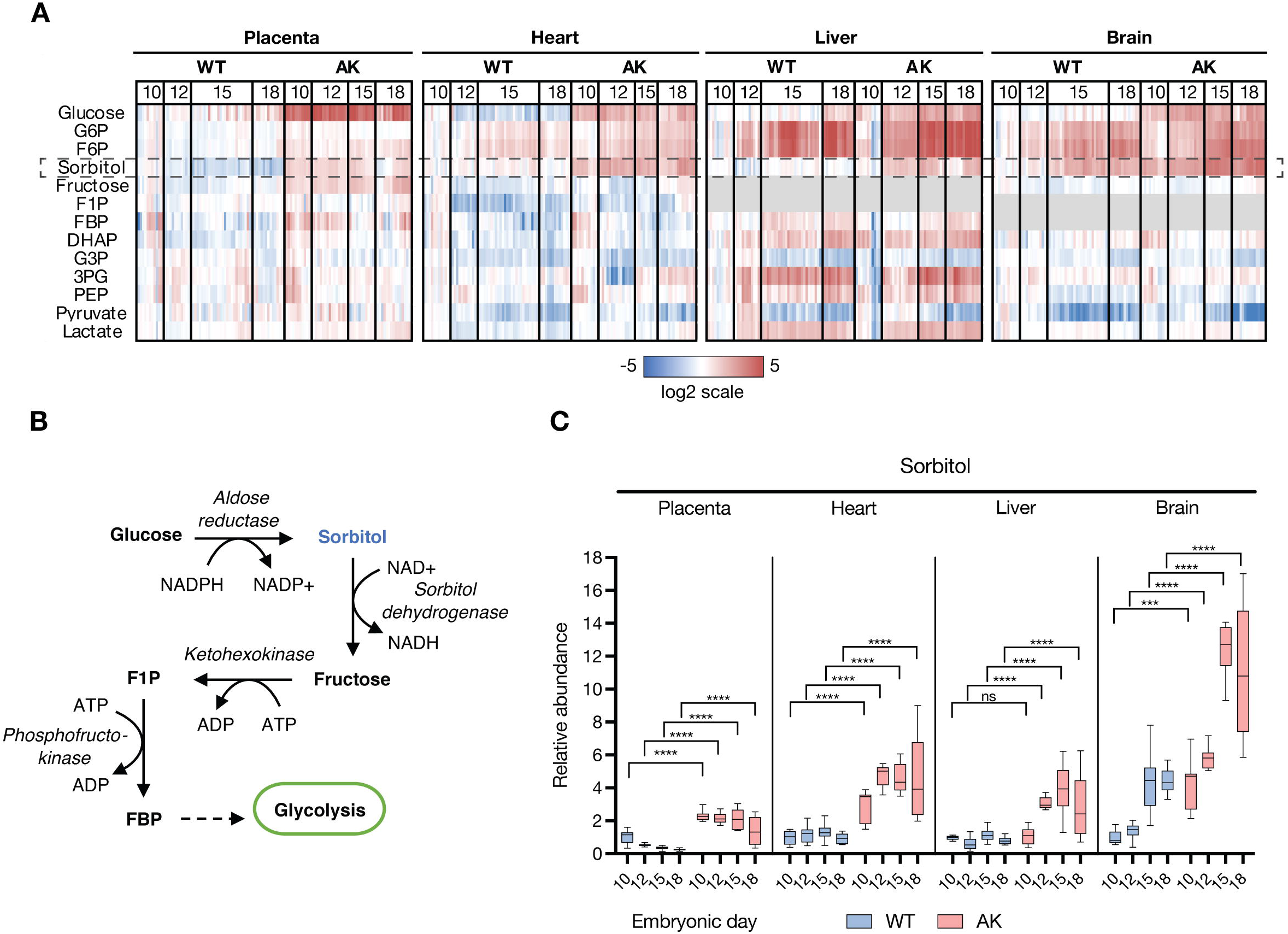
Maternal hyperglycemia causes sorbitol accumulation in fetal tissues. (A) Heatmap representing the log2 fold changes in glycolytic and polyol pathway intermediates relative to E10.5 fetal tissues from wildtype (WT) dams. Gray indicates not detected. (B) Schematic of the polyol pathway and its intersection with glycolysis. (C) Sorbitol pool levels in fetal tissues isolated from WT and AK dams. Data represented as a relative comparison to E10.5 fetal tissues from WT dams. Individual values are plotted relative to the mean of WT E10.5. Tukey method was applied for plotting whiskers and outliers. Statistical analyses were performed using two-way ANOVA.

### Maternal hyperglycemia alters amino acid metabolism in the fetal brain

We were curious how maternal hyperglycemia altered broader fetal metabolism. We observed changes in amino acid levels in fetal tissues harvested from wildtype versus Akita dams (Figure 3A). Over the developmental stages, most amino acid levels in fetal hearts harvested from Akita dams decreased to a lesser extent compared to fetal hearts harvested from wildtype dams, while those in Akita fetal livers and brains trended up more strongly. Most notably, fetal brain aspartate levels presented different trends over the developmental timeframe assessed in fetuses harvested from wildtype versus Akita dams (Figure 3B). In fetal brains isolated from wildtype dams, aspartate levels increased roughly 2-fold from E10.5 to E12.5, however in fetal brains isolated from Akita dams, aspartate levels remained unchanged from E10.5 to E12.5 and instead increased roughly 2-fold between E12.5 to E15.5 (Figure 3B). These changes are in contrast to fetal brain asparagine levels which remained unchanged across the developmental timeframe and conditions assessed (Figure 3B). The levels of glutamate and glutamate-derived gamma-aminobutyric acid (GABA) were also different in fetal brains harvested from wildtype versus Akita dams (Figure 3C). Glutamate and GABA levels were lower in E10.5 and E12.5 fetal brains isolated from Akita dams relative to E10.5 and E12.5 fetal brains isolated from wildtype dams. Lower levels of these neurotransmitters may contribute to the higher incidence of congenital brain defects observed in fetuses exposed to maternal hyperglycemia (Represa and Ben-Ari, 2005).

**Figure 3.**
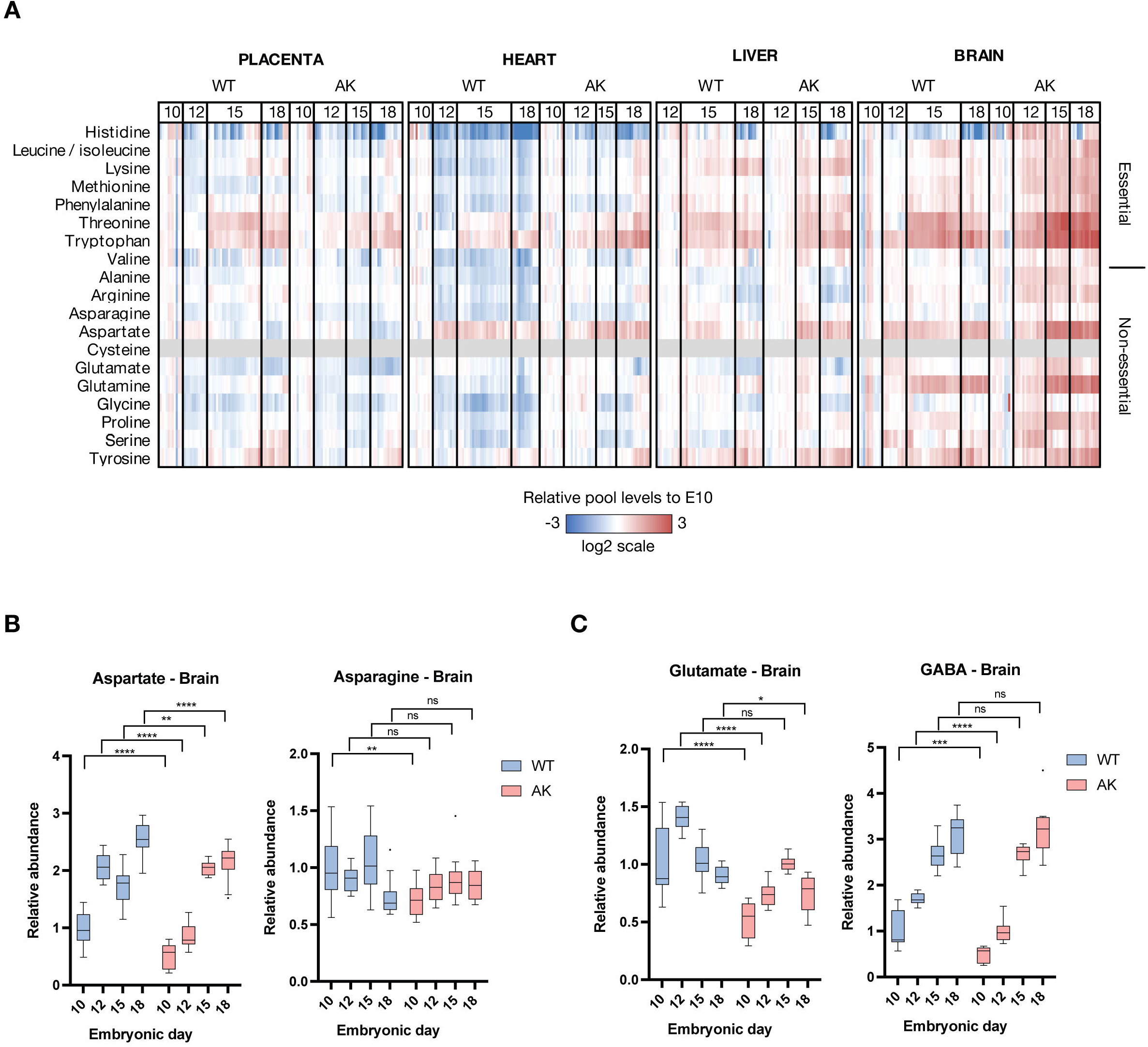
Maternal hyperglycemia alters aspartate and glutamate levels in fetal brains during mid-to-late gestation. (A) Heatmap representing the log2 fold changes in amino acid pool levels relative to E10.5 fetal tissues isolated from both wildtype (WT) and Akita (AK) dams. For fetal tissues from WT dams, data is represented relative to the WT E10.5 samples. For fetal tissues from AK dams, data is represented relative to the average of AK E10.5 samples (B) Relative aspartate and asparagine levels in fetal brain tissues. Data represented relative to E10.5 fetal brain tissues from WT dams. Tukey method was applied for plotting whiskers and outliers. Statistical analyses were performed using two-way ANOVA. (C) Relative glutamate and GABA levels in fetal brain tissues. Data represented relative to E10.5 fetal brain tissues from WT dams. Tukey method was applied for plotting whiskers and outliers. Statistical analyses were performed using two-way ANOVA.

### Isotope tracing reveals fetal growth strategies

Tracing [U-^13^C]glucose past central carbon metabolism revealed how fetal tissues source biomass building blocks. Using the principle that a metabolite’s ^13^C labeling fraction cannot be greater than that of its ^13^C source, we traced ^13^C at two different levels: 1) the wholebody level, which reveals nutrient exchange fluxes, and 2) the tissue level, which reveals fluxes through metabolic pathways. At the whole-body level, we considered the fetal circulatory system to be connected to maternal circulation through a path from the placenta to the liver, to the heart, and to the brain (Kiserud, 2005;Linask et al., 2014) (Figure 4A). We observed that seven non-essential amino acids (Asp, Asn, Glu, Gln, Pro, Ser, and Gly) were ^13^C-labeled in fetuses, and of those seven, aspartate and glycine not labeled in maternal plasma (Figure 4B). Therefore, aspartate and glycine were entirely synthesized by fetal tissues. Maternal and fetal metabolism contributed to five amino acids (Asn, Glu, Gln, Pro, and Ser) in fetuses, and the 13 other amino acids were entirely sourced from the dam. The uptrend of glutamate labeling through the circulatory path in E15.5 and E18.5 fetuses implied that every fetal tissue made its own glutamate, whereas the serine labeling peaking in the liver implied that the heart and the brain may not necessarily make their own serine and depend on serine overflow from the liver. Similarly, proline in the placenta may be contributing to fetal tissues in E10.5 before the liver and the brain develops the proline synthesis capability as evidenced in E15.5 fetuses. The time development of asparagine labeling revealed that asparagine biosynthesis in fetal livers isolated from wildtype dams begins at E18.5 while it begins earlier at E15.5 in fetal livers isolated from Akita dams (Figure S3A).

**Figure 4.**
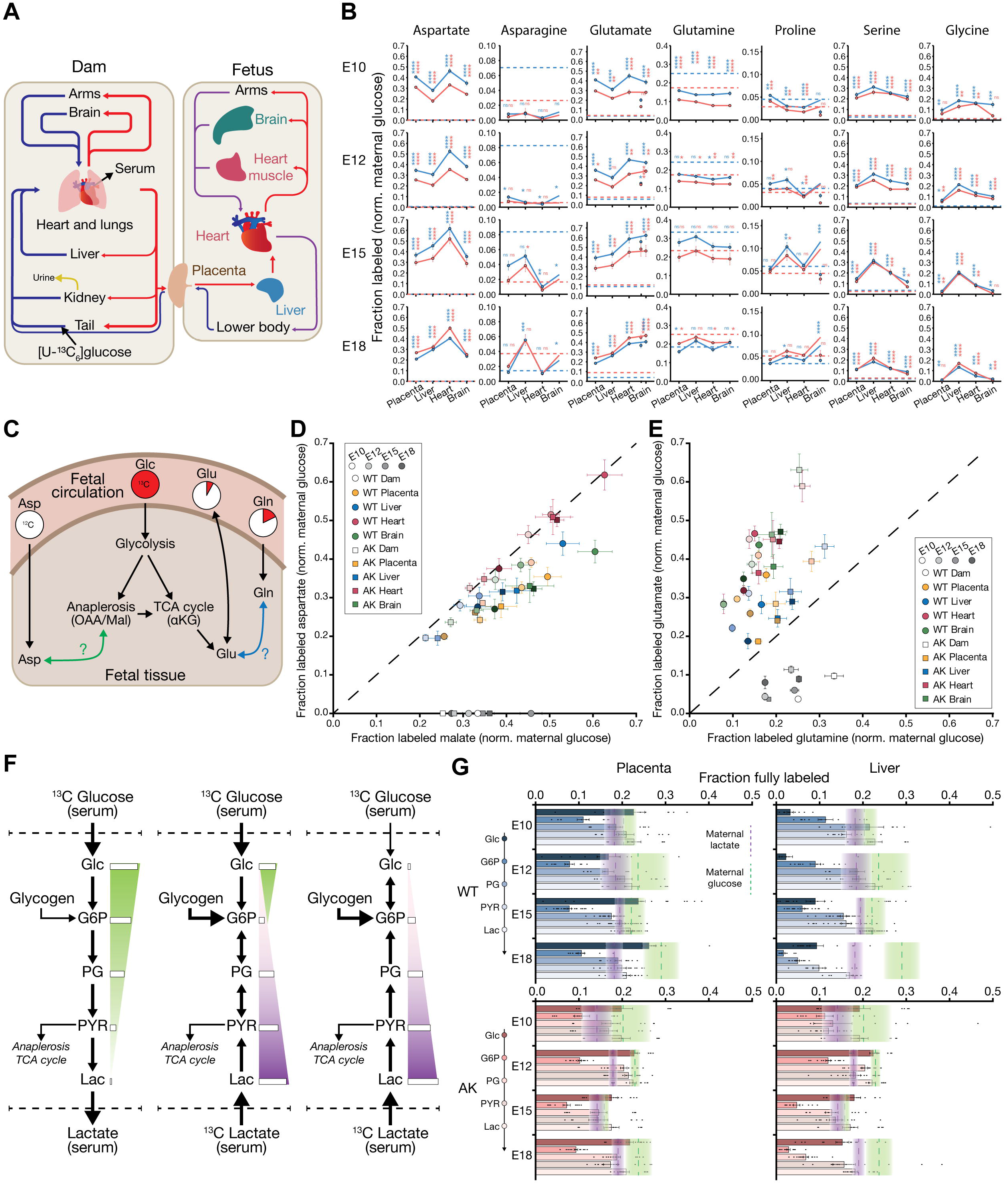
Isotope tracing in fetal tissues elucidates fetal nutrient sourcing and utilization. (A) Schematic of fetal circulation from the dam to the fetus. Nutrients from the dam are transferred through the placenta and into fetal circulation, where the liver, heart, and brain can uptake nutrients supplied via the dam. (B) Tracing of seven amino acids, which were labeled in fetal tissues, through fetal circulation. Labeling across four fetal tissues (placenta, liver, heart, brain) was compared to the labeling of circulating amino acids in the maternal bloodstream (dashed line). Blue lines indicate amino acid labeling in fetal tissues from WT dams, and red lines indicate labeling in fetal tissues from Akita dams. Statistics between the tissue and dam were performed under a two-tailed *t*-test. (C) Schematic of the sources of non-essential amino acids. (D) Labeling of aspartate and malate across fetal tissue and maternal plasma. The dashed line represents the line of unity, where malate and aspartate labeling are equal. (E) Labeling of glutamate and glutamine across fetal tissue and maternal plasma. (F) Depiction of three major sources of carbon for glycolytic intermediates and biomass building blocks. Glycolysis and gluconeogenesis both can supply ^13^C to glycolytic intermediates. The labeling of glycolytic intermediates may follow labeling of either glucose (green) or lactate (purple). Glycogen, which is unlabeled, may dilute labeling of glycolytic intermediates. (G) Fraction of fully labeled glycolytic intermediates (glucose, glucose-6-phosphate, phosphoglycerate, pyruvate, lactate) in WT and AK placenta and liver compared to maternal glucose and lactate labeling. Green bands represent maternal plasma labeling of glucose, and purple bands represent maternal labeling of lactate. Mean ± s.e.m. are represented by a dashed line and the width of the bands.

At the tissue level, ^13^C tracing revealed the metabolic pathways responsible for the labeling of amino acids (Figure 4C). Tissue metabolites can gain ^13^C via maternofetal [U-^13^C]glucose transport through the placenta and via transport of other circulating metabolites the dam had produced from [U-^13^C]glucose (Baumann et al., 2002;Lager and Powell, 2012). Fetal aspartate was similarly or less labeled than malate, a proxy for oxaloacetate that is the direct precursor of aspartate, but more labeled than plasma aspartate (Figure 4D). Fetal glutamate labeling was consistently higher than both plasma glutamate labeling and fetal glutamine labeling but comparable to tissue α-ketoglutarate labeling (Figure 4E and S3B). These observations implied that fetal tissues *de novo* synthesized aspartate and glutamate via anaplerosis and the TCA cycle.

We set out to determine if the ^13^C labeling in fetal tissues was directly or indirectly from glucose. Since circulating lactate is another high-flux carbon source, we considered three scenarios that yield different ^13^C labeling: the major carbon source(s) being 1) plasma glucose, 2) both plasma glucose and lactate, and 3) plasma lactate (Figure 4F and S3C) (Hui et al.,2017). Additionally, we considered the potential introduction of unlabeled hexose phosphate via glycogenolysis (TeSlaa et al., 2021). In the placenta, glycolytic labeling reflected the contribution of both plasma glucose and plasma lactate, which had been labeled in the dam across mid-to-late gestation (Figure 4G, S3D, and S3E). The liver, on the other hand, displayed disparate glycolytic labeling signatures between fetuses from wildtype and Akita dams. In E10.5 and E12.5, fetal liver from wildtype dams displayed ascending labeling gradient across glycolysis, indicating gluconeogenesis from ^13^C-labeled lactate and perhaps dilution of hexose ^13^C labeling by glycogenolysis (Figure S3F). However, ^13^C glucose labeling was higher in fetal livers harvested from Akita dams across mid-to-late gestation and more similar to the level of glucose labeling in maternal circulation. Taken together, maternal hyperglycemia not only alters the source of carbon backbones in fetuses but also sustains glucose contribution to amino acid biosynthesis in late gestation.

### Fetal nucleotide synthesis slows in mid-to-late gestation

Nucleotides are in high demand during rapid cell division, yet they are not available via circulation. Thus, we investigated how fetal tissues manage nucleotide biosynthesis. We observed fetal tissue-specific increases in nucleotides in the heart, liver, and brain - but not placenta - at E12.5, E15.5, and E18.5 relative to E10.5 (Figure 5A and S4A). Nucleoside monophosphates and diphosphates, but not nucleoside triphosphates, are elevated in fetal brain tissues at E15.5 and E18.5 relative to E10.5. However, almost all nucleotides measured are elevated in E12.5-E18.5 fetal liver tissues relative to E10.5. Specific nucleotides, such as IMP, ADP and ATP, but not AMP, CMP, GMP or UMP, were most elevated in fetal heart tissues at E15.5 and E18.5 relative to E10.5. We were interested to find how purine and pyrimidine metabolism changed across fetal tissue throughout mid-to-late gestation. In placenta and fetal heart, AMP and UMP pool levels did not change at the developmental stages we measured (Figure 5B and 5C). However, in fetal liver and brain, AMP and UMP pool levels increased (Figure 5B and 5C). Additionally,^13^C enrichment into AMP and UMP consistently decreased across fetal tissues as embryonic day advanced, regardless of pool level changes (Figure 5D and 5E). Similarly, decreasing labeling was observed for IMP and GMP irrespective of pool level trends (Figure S4B-G).

**Figure 5.**
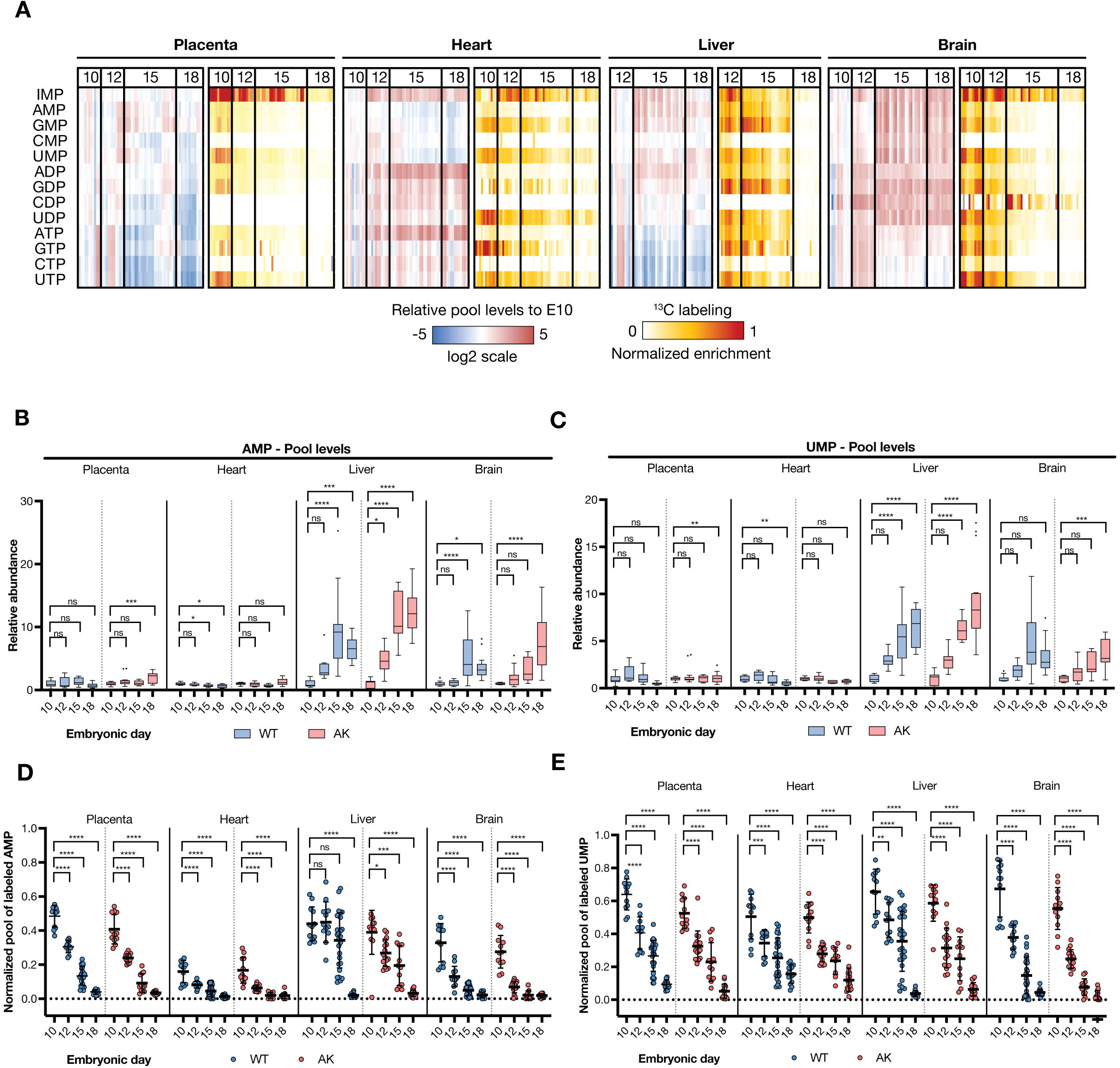
Progressive decrease in *de novo* nucleotide synthesis during fetal development. (A) Blue to red scale: Heatmap representing the log2 fold changes in nucleotide pool levels relative to E10.5 fetal tissues from WT dams. White to red scale: Heatmap representing the ^13^C-labeled pool of nucleotides in fetal tissues from WT dams. (B) Relative AMP pool levels across fetal tissues. For fetal tissues from WT dams, data is represented relative to the average of WT E10.5 samples. For fetal tissues from Akita (AK) dams, data is represented relative to the average of AK E10.5 samples. Tukey method was applied for plotting whiskers and outliers. Statistical analyses were performed using two-way ANOVA. (C) Relative UMP pool levels across fetal tissues. For fetal tissues from WT dams, data is represented relative to the average of WT E10.5 samples. For fetal tissues from Akita (AK) dams, data is represented relative to the average of AK E10.5 samples. Tukey method was applied for plotting whiskers and outliers. Statistical analyses were performed using two-way ANOVA. (D) Normalized pool of ^13^C-labeled AMP across fetal tissues. For each fetal tissue sample, the AMP labeled pool was normalized to its respective glucose labeled pool from maternal plasma. (E) Normalized pool of labeled of UMP across fetal tissues. For each fetal tissue sample, the UMP labeled pool was normalized to its respective glucose labeled pool from maternal plasma.

Despite the uniform ^13^C-glucose infusion procedure, ^13^C enrichment in nucleotides decreased from mid-to late-gestation (Figure 5A and S4A). Nucleotide labeling, which did not reach isotopic steady state in three hours, indicated nucleotide turnover rates. Thus, the decreased labeling that coincided with level or decreased pool sizes implied slower nucleotide biosynthesis fluxes. We observed that nucleotide biosynthesis in the placenta and the heart started slowing down in earlier developmental stages (i.e., E10.5 or E12.5) than in the liver and the brain (i.e., E15.5 or later).

### Histidine-derived metabolites accumulate in late-term fetuses

We expanded our analysis platform to survey all detected LC-MS peaks in an untargeted manner and identify metabolic signatures that characterize developmental transitions. We compared untargeted data from E18.5 and E10.5 placentas harvested from both wildtype and Akita dams (Table S3) using the MetaboAnalyst platform (Pang et al., 2022). This analysis yielded histidine metabolism as the metabolic pathway with the highest normalized enrichment score and significance in E18.5 versus E10.5 placentas harvested from wildtype dams (Figure 6A). Similar results were found when comparing placental metabolites from fetuses harvested from Akita dams (Figure S5A). Given the significant enrichment of histidine metabolism in late stage versus E10.5 placentas, we decided to look more closely at this pathway across fetal tissues and throughout mid-to-late gestation. Histamine is derived from histidine, which can be catabolized to urocanate as well as to histamine (Figure 6B). We found that histidine pool levels did not follow a generalized trend across fetal tissues, and in fact, decreased in fetal hearts and brains isolated from wildtype dams (Figure 6C). However, we observed a stark accumulation of histidine-derived metabolites at the E18.5 stage in fetal placenta, heart, liver, and brain tissues (Figure 6D and 6E). In particular, we observed accumulation of the histidine degradation products, urocanate, as well as histamine and imidazole-4-acetate (Figure 6D-F). These identified signals through untargeted analysis were corroborated by chemical standards (Figure S5B). The increase in histidine-derived metabolites at E18.5 was also observed in maternal circulation (Figure 6G), underscoring the interaction between maternal and fetal metabolism. Intriguingly, elevated histamine plasma levels have been detected in women during preterm labor compared with term labor (Maintz et al., 2008).

**Figure 6.**
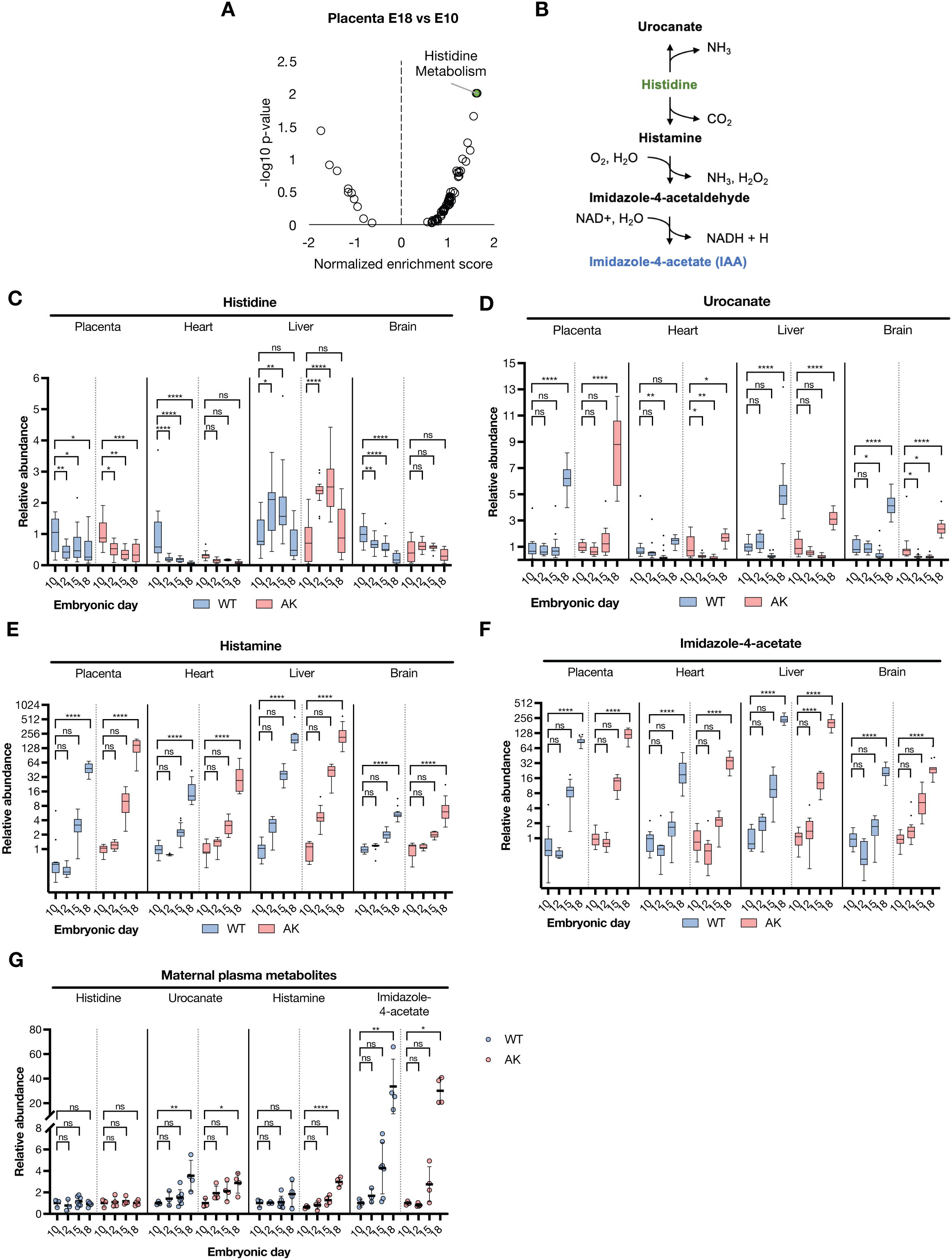
Histidine-derived metabolites accumulate in fetal tissues during late development. (A) MetaboAnalyst pathway analysis using positive ion mode untargeted data highlights at changes in histidine metabolism in placenta E18.5 tissues relative to E10.5 from wildtype dams. (B) Histidine metabolism pathway schematic. (C-F): Relative pool levels of histidine (C) and its derived metabolites, urocanate (D), histamine (E), and imidazole-4-acetate (F) across fetal tissues from both wildtype (WT) and Akita (AK) dams. For fetal tissues from WT dams, data is represented relative to the average of WT E10.5 samples. For fetal tissues from Akita (AK) dams, data is represented relative to the mean of AK E10.5 samples. Tukey method was applied for plotting whiskers and outliers. Statistical analyses were performed using twoway ANOVA. Relative pool levels of histamine (E) and imidazole-4-acetate (F) are shown using a log2 scale for the y-axis. (G) Relative pool levels of histidine derived metabolites in maternal plasma. For maternal plasma from WT dams, data is represented relative to the mean of WT E10.5 samples. For maternal plasma from Akita (AK) dams, data is represented relative to the mean of AK E10.5 samples. Error bars denote standard deviation. Statistical analyses were performed using two-way ANOVA.

## DISCUSSION

Our extensive metabolomic dataset presents the first in-depth assessment of *in vivo* fetal tissue metabolism during mid-to-late gestation within the context of both maternal euglycemia and hyperglycemia. Examination of relative metabolite pool levels in fetal tissues revealed elevated levels of the glucose-derived toxic metabolite sorbitol in all fetal tissues examined from hyperglycemic dams (Figure 2). These findings suggest that fetal tissues are not shielded from sorbitol accumulation when developing under maternal hyperglycemia. Uncontrolled high blood glucose in diabetic patients is recognized to lead to sorbitol accumulation and tissue damage in insulin-independent tissues such as retina, kidney, and nerves (Brownlee, 2001). Early research hinted at sorbitol accumulation in fetuses from diabetic rats (Eriksson et al., 1986). Our *in vivo* metabolomics data confirms that fetuses from diabetic mice exhibit 3-to 5-fold sorbitol accumulation compared to wildtype. Further studies should interrogate the impact this amount of sorbitol accumulation has on fetal development. Inhibitors of the sorbitol producing enzyme, aldose reductase, have been used in the clinic to treat diabetic neuropathy (Schemmel et al.,2010). Past research using aldose reductase inhibitors on cultured rat conceptuses resulted in reduction of sorbitol with no effect on prevention of high glucose-induced growth retardation and dysmorphogenesis (Hod et al., 1986). However, more research will be needed to test whether aldose reductase inhibitors can be used to reduce fetal sorbitol levels *in vivo* and if this prevents of some of the developmental defects associated with maternal hyperglycemia.

Additionally, our analysis of relative metabolite pool levels in fetal tissues during mid-to-late gestation revealed altered trends in fetal brain amino acid levels (Figure 3). While each analyzed tissue presented unique trends in amino acid levels throughout mid-to-late gestation, we highlighted those of the brain given the important role certain amino acids have as neurotransmitters in shaping brain networks (Kölker, 2018). For instance, while there is growing appreciation on the importance of the glutamatergic system in brain development and function (Matsugami et al., 2006), there is a dearth of information on how metabolic abnormalities impact such systems. Therefore, our findings that amino acid neurotransmitters such as aspartate, glutamate, and GABA exhibit different levels in fetal brains isolated from hyperglycemic dams suggest follow-up studies examining whether these alterations may contribute to the higher incidence of congenital brain defects occurring in the context of maternal hyperglycemia.

Beyond understanding metabolite pool level dynamics, we leveraged ^13^C-labeling information to investigate fetal tissue central carbon pathway usage as well as nutrient sourcing. For instance, we found that differences in fetal and plasma nutrient labeling from the ^13^C-glucose infused into the dam can inform whether a particular nutrient is supplied via maternal circulation or whether it is synthesized by the fetus. Through this analysis, we found that fetal tissues exhibit aspartate and glutamate synthesis throughout mid-to-late gestation, however proline biosynthesis becomes more prominent in late liver and brain development. Additionally, our data suggest that fetal liver tissue derives carbon backbones from a different source than circulating glucose, but less so in the case of maternal hyperglycemia, in which fetuses sustained glucose dependence. This raises an important consideration of carbon sourcing in the developing fetus, particularly at E10.5. In euglycemia, fetuses obtain carbon backbones flexibly via glycogen breakdown or from lactate via gluconeogenesis. Complementing our studies presented here with additional labeling experiments that incorporate other labeled nutrients such as ^13^C-lactate represents a logical future direction, which will help illuminate the alternative carbon sources that are relied on by E10.5 livers and hearts from euglycemic and hyperglycemic dams. Notwithstanding, this interesting shift in carbon sourcing in fetuses from hyperglycemic dams highlights the metabolic plasticity and adaptations fetal tissues undergo when challenged with a changing nutrient microenvironment.

Untargeted analyses of our large dataset revealed the striking accumulation of histidine-derived metabolites, including histamine and imidazole-4-acetate, in fetal tissues and maternal plasma at E18.5 (Figure 6). Intriguingly, elevated histamine plasma levels have been detected in women during preterm compared with term labor (Maintz et al., 2008). These metabolites are therefore of special interest given their potential role in promoting labor. The stark increase of histidine-derived metabolites we observed as fetuses enter the late stage E18.5 underscores how little is known about the potential role metabolism has in modulating developmental transitions.

Overall, the data resource presented here will help initiate new hypotheses for future testing that will lead to more mechanistic insights connecting metabolism to cell fate transitions. For example, future studies should examine whether metabolites we found elevated in the fetal tissues during diabetic pregnancy are responsible for the increased incidence of congenital defects, as well as determine if sharp metabolite fluctuations delineate important developmental transitions. Better understanding of how maternal metabolism impacts fetal metabolism and development will be crucial to design strategies that promote maternal health and address the urgent clinical issue of diabetic pregnancy.

## LIMITATIONS OF THE STUDY

Although mouse development does not entirely mimic human development, we expect our metabolomic dataset to serve as a valuable complement to extensive studies looking at development through a genomics and transcriptomics lens. Due to the nature of our experiments that necessitated careful timing of pregnancies, we decided to streamline our platform to implement tail vein infusions, which constrained our ability to perform longer infusions and therefore limited the number of metabolites that incorporate the ^13^C-label from [U-^13^C]glucose. Given the number of mice infused in this study and the number of fetuses collected, we centered our experimental approach around using a single isotopically labeled nutrient, [U-^13^C]glucose, and the collection of four tissues per fetus dissected. Utilization of additional tracers such as ^13^C-glutamine and ^13^C-lactate in future studies, will provide additional layers with which to assess pathway use. Testing of additional embryonic days beyond the 4 representative stages would provide a more precise dissection of fetal metabolic networks. Despite sample normalization, we note that E10.5 fetal livers presented significantly lower metabolite pool levels across many metabolites when compared to E12.5 samples. While this difference was not present among many metabolites, we acknowledge it may be unclear whether this trend has biological value or rather reflects a potential underrepresentation of metabolite pool levels. We recognize that use of the Akita mouse model is one several approaches that are used to model diabetes in the mouse. While the Akita mouse does not perfectly represent all the mechanistic features that may characterize human diabetic pregnancies, our goal with this study is to provide a starting point to analyze the impact high glucose *in utero* has on the metabolism of the developing fetus.

## AUTHOR CONTRIBUTIONS

HN, AN, and HRC conceptualized the examination of metabolic time course of developing organs. CAP, HN, AN, and HRC designed the study. CAP optimized and performed the *in vivo* [U-^13^C]glucose infusions, conducted all fetal tissue and plasma metabolite extractions, and performed the labeling analysis and targeted/untargeted metabolomics data analysis. HN set up timed pregnancies, collected blood and measured maternal glucose levels, counted fetal defects, and dissected fetal tissues. NM ran the LC-MS samples and performed raw file processing and targeted peak assignment. RCL and JOP analyzed the labeling data and developed models to assess nutrient sourcing and pathway utilization. JT assisted with fetal tissue processing and AP assisted with *in vivo* [U-13C]glucose infusions and metabolite extractions. CAP, HN, RCL, JOP, AN, and HRC conducted data interpretation and wrote and/or edited original manuscript with input from all authors.

## METHODS

### Lead contact and materials availability

Further information and requests for resources and reagents should be directed to and will be fulfilled by the Lead Contact, Heather R. Christofk.

### Data availability

The metabolomics data reported in this study will be deposited in the National Metabolomics Data Repository (NMDR). To request access, contact Heather R. Christofk.

### Experimental model and subject details

#### Mice

Mice were housed in pathogen-free facilities at University of California Los Angeles (UCLA). All animal experiments were approved by the UCLA Animal Research Committee (ARC), and we complied with all relevant ethical regulations while conducting animal experiments.

### Method details

#### Animal studies

Healthy wildtype and Akita dams were set up for mating. The following morning, females displaying vaginal plugs were identified as pregnant, recorded as embryonic day (E) 0.5 and moved to a new cage until the appropriate embryonic day to be interrogated.

#### Pregnant mouse infusions

Labeled glucose solution was prepared at a concentration of 100 mg/mL in filtered 0.9% sodium chloride solution. After overnight fasting (from 18:00 the day before), infusions took place around between 09:00 and 10:00 for all pregnant dams. Mice were anesthetized using isoflurane gas at 5% and placed on a warm pad. Mice were then kept under 2.5% isoflurane for the duration of infusion. For catheter placement, a 28-gauge insulin syringe needle was connected via polyethylene tubing (PE-10) to a syringe (containing glucose solution) placed on an infusion pump (Harvard Apparatus). At the start of the infusion, the 28-gauge needle was inserted into the tail vein. A glucose bolus of 4 μL/gBW was administered. Right after bolus administration, infusion rate was set at a continuous 0.085ul/gBW for a total infusion time of 3 hours.

#### Fetal tissue extraction

Following infusion, mice were euthanized and blood was collected via heart puncture. Fetal tissues (placenta, brain, liver, and heart) were dissected in ice-cold sterile PBS. Immediately after dissection, weight was recorded and fetal tissue was placed in a pre-filled bead mill tube containing metal beads and 500 μL of methanol:water (80:20) solution kept cold on dry ice. Fetal tissues were homogenized using a Fisherbrand™ Bead Mill Homogenizer. Samples were spun at 17,000 g (4 °C) for ten minutes to remove precipitated cell material (protein/DNA). Supernatants were collected, transferred to a clean tube, and evaporated using a Nitrogen evaporator (Organomation). Evaporated metabolite extracts were stored at −80 °C. Pellets containing protein/DNA were dried on a heat block (55 °C) and stored at −80 °C

#### Maternal plasma extraction

Collected blood was centrifuged at 5,000g to collect plasma. Plasma was snap frozen in liquid nitrogen and stored until extraction. For metabolite extraction 5 μL plasma was mixed with 500 μL methanol:water (80:20) solution (−80 °C). Samples were centrifuged for ten minutes at 17,000 *g* (4 °C) and 450 μL of each sample was evaporated using a Nitrogen evaporator (Organomation). Evaporated metabolite extracts were stored at −80 °C.

#### Fetal tissue DNA measurements and weight normalization

Frozen pellets were resuspended in a solution containing 100mM NaCl, 20mM Tris-HCl (pH 7.4), 5mM EDTA, 0.1% SDS, Proteinase K (500 μg/mL). Volume for pellet resuspension varied per size of fetal tissue pellet to ensure all material was dissolved. Resuspension volume was recorded as a dilution factor. DNA concentration was measured using a nanodrop and total DNA concentration was calculated via dilution factor (ThermoFisher Scientific).

#### Metabolite measurement by LC-MS

Dried metabolite extracts were reconstituted in 50% acetonitrile (ACN) 50%dH20 solution. For fetal liver, brain, and placentas, metabolite extract resuspension volume was set to 75 μL per mg weight. For fetal hearts, metabolite extract resuspension volume was set to 90 μL per mg weight. To increase accuracy in metabolite resuspension volumes, total DNA concentration measurements from each fetal tissue group were correlated to directly measured tissue weights higher than 2mg, and a linear regression equation was used to calculate weight values for tissue samples with recorded weights below 2mg. For plasma metabolite extracts, resuspension volume was set to 100 μL. After resuspension, samples were vortexed and spun down for 10 min at 17,000g. 75 μL of the supernatant was then transferred to HPLC glass vials. Samples were run on a Vanquish (Thermo Scientific) UHPLC system with mobile phase A (20mM ammonium carbonate, pH 9.7) and mobile phase B (100% ACN) at a flow rate of 150 μL/min on a SeQuant ZIC-pHILIC Polymeric column (2.1 × 150 mm 5 μm, EMD Millipore) at 35°C. Injection volume was set to 10 μL. Separation was achieved with a linear gradient from 20% A to 80% A in 20 min followed by a linear gradient from 80% A to 20% A from 20 min to 20.5 min. 20% A was then held from 20.5 min to 28 min. The UHPLC was coupled to a Q-Exactive (Thermo Scientific) mass analyzer running in polarity switching mode with spray-voltage=3.2kV, sheath-gas=40, aux-gas=15, sweep-gas=1, aux-gas-temp=350°C, and capillary-temp=275°C. For both polarities mass scan settings were kept at full-scan-range=(70-1000), ms1-resolution=70,000, max-injection-time=250ms, and AGC-target=1E6. MS2 data was also collected from the top three most abundant singly charged ions in each scan with normalized-collision-energy=35. Each resulting .raw files was centroided and converted into two .mzXML files (one for positive ion mode scans and one for negative ion mode scans) using msconvert from ProteoWizard (Chambers et al., 2012).

#### Metabolomic data analysis

.mzXML files were imported into the MZmine 2 software package (Pluskal et al., 2010). Ion chromatograms were generated from MS1 spectra via the built-in Automated Data Analysis Pipeline (ADAP) (Myers et al., 2017). Chromatogram module and peaks were detected via the ADAP wavelets algorithm. Peaks were aligned across all samples via the Random sample consensus aligner module, gap-filled, and assigned identities using an exact mass MS1(±15ppm) and retention time RT (±0.5min) search of our in-house MS1-RT database. Peak boundaries and identifications were then further refined by manual curation. Peaks were quantified by area under the curve integration and exported as .CSV files. For isotopologue analysis, peak areas were processed via the R package AccuCor to correct for natural isotope abundance (Su et al., 2017). Peak selection for untargeted metabolomic analysis was performed using El-Maven (Elucidata) automatic feature detection. Functional analysis annotation of untargeted data was performed using MetaboAnalyst 5.0, using peak intensity tables as input, with a mass tolerance set at 15 ppm, and processing the data through an interquartile range (IQR) filter, a log base 10 transformation, and the automatic data scaling feature. (Pang et al., 2022). Gene set enrichment analysis algorithm, integrated in the MetaboAnalyst pipeline was applied using the KEGG *(mus musculus)* database as reference.

## SUPPLEMENTAL FIGURE LEGENDS

**Figure S1.**
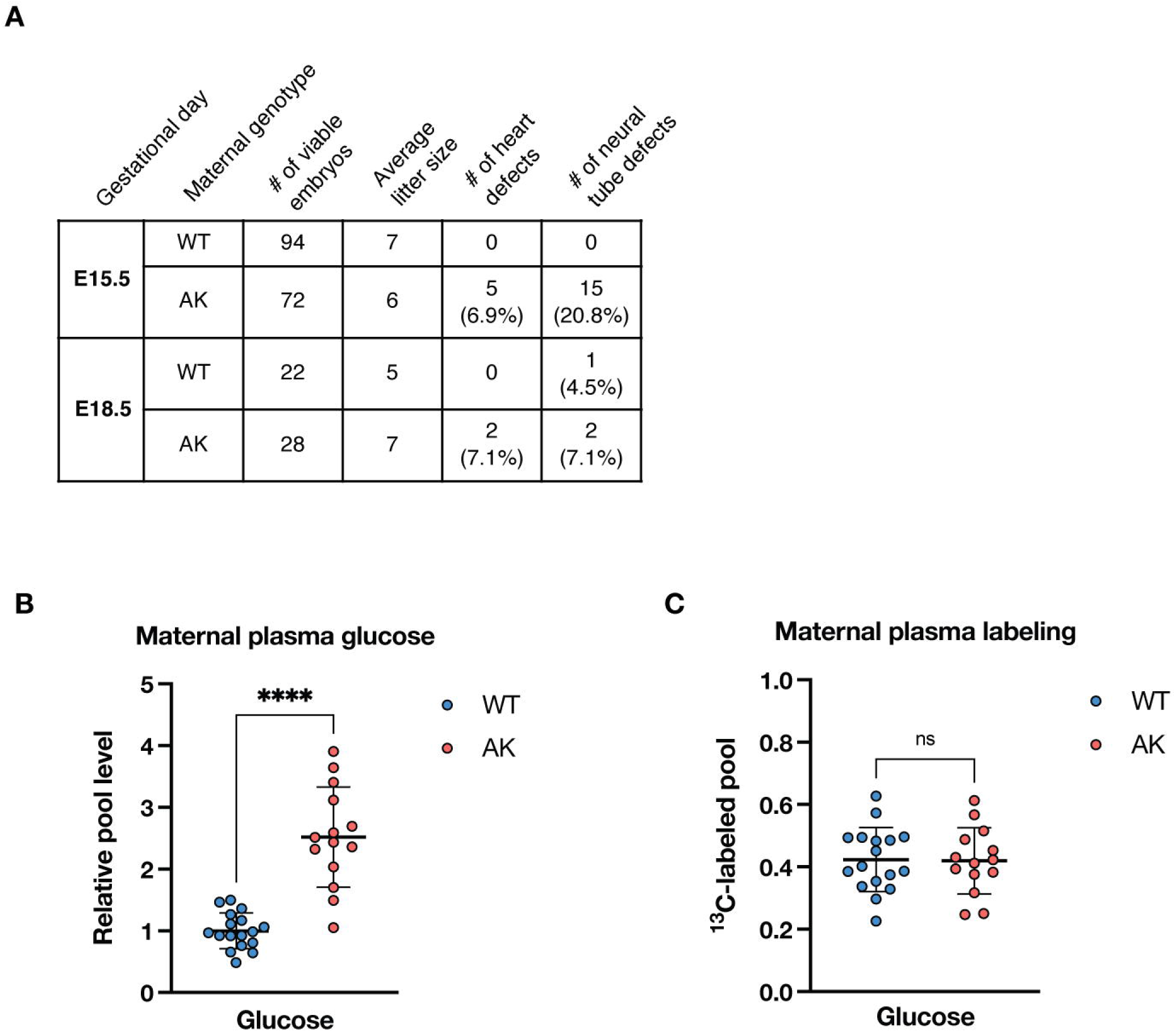
Use of pregnant Akita mice to model maternal hyperglycemia, related to Figure 1. (A) Table describing number of heart and neural defects observed in fetuses from a representative cohort of wildtype (WT) versus Akita (AK) dams. Only fetuses lacking severe defects were used for this study. (B) Relative plasma glucose levels in the pregnant wildtype dams relative to Akita dams used in this study. Error bars denote standard deviation. Statistical analyses were performed using two-tailed *t*-tests. (C) Levels of ^13^C glucose enrichment in the plasma of the pregnant wildtype dams relative to Akita dams used in this study. Error bars denote standard deviation. Statistical analyses were performed using two-tailed *t*-test.

**Figure S2.**
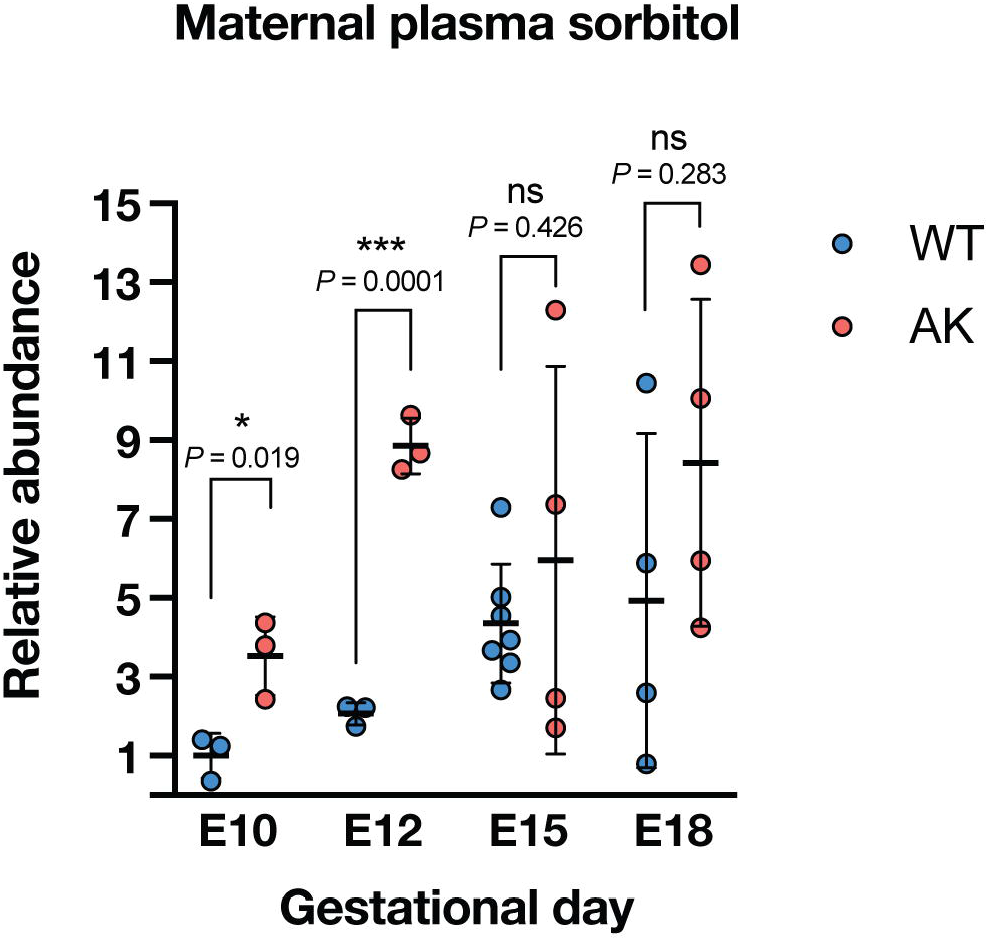
Elevated sorbitol levels in E10.5 and E12.5 Akita dams, related to Figure 2. Relative pool levels of sorbitol in maternal plasma. Data is represented relative to the mean of WT E10.5 samples. Error bars denote standard deviation. Statistical significance was determined using two-tailed *t*-tests.

**Figure S3.**
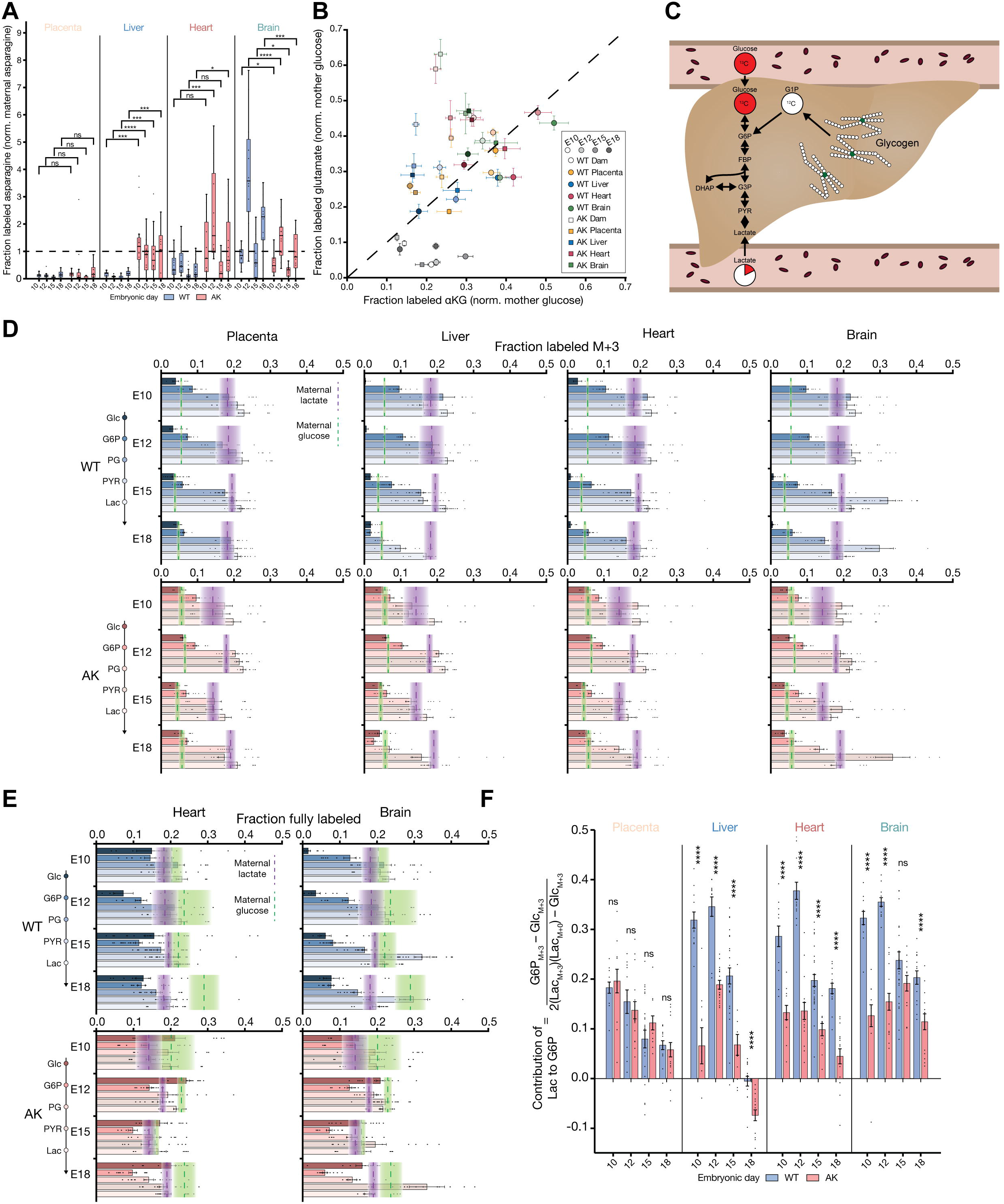
Isotope tracing in fetal tissue elucidates metabolic strategy, related to Figure 4. (A) Labeling of fetal asparagine normalized to maternal asparagine labeling in wildtype (WT) and Akita (AK) dams. Statistical analyses were determined using two-tailed *t*-tests. (B) Labeling of intracellular glutamate (vertical) compared to intracellular α-ketoglutarate (horizontal). The dashed line represents the line of parity, where glutamate and α-ketoglutarate labeling are equal. (C) Schematic of glycogen breakdown relative to glycolysis. Glycogen, which has slow turnover relative to glucose, introduces unlabeled hexose phosphates into glycolysis. (D) M+3 labeling of glycolytic intermediates from glucose to lactate in fetal tissues. Green bands represent maternal plasma labeling of glucose and purple bands represent maternal labeling of lactate. Means ± s.e.m. are represented by a dashed line and the width of the bands. (E) Fraction of fully labeled glycolytic intermediates (glucose, glucose-6-phosphate, phosphoglycerate, pyruvate, lactate) in fetal hearts and brains from WT and AK dams compared to maternal glucose and lactate labeling. Green bands represent maternal plasma labeling of glucose, and purple bands represent maternal labeling of lactate. Means ± s.e.m. are represented by a dashed line and the width of the bands. (F) Comparison of lactate labeling contribution to G6P in WT and AK tissues. Negative values indicate substantial glycogenolysis. Statistical analyses between WT and AK were determined using two-tailed *t*-tests.

**Figure S4.**
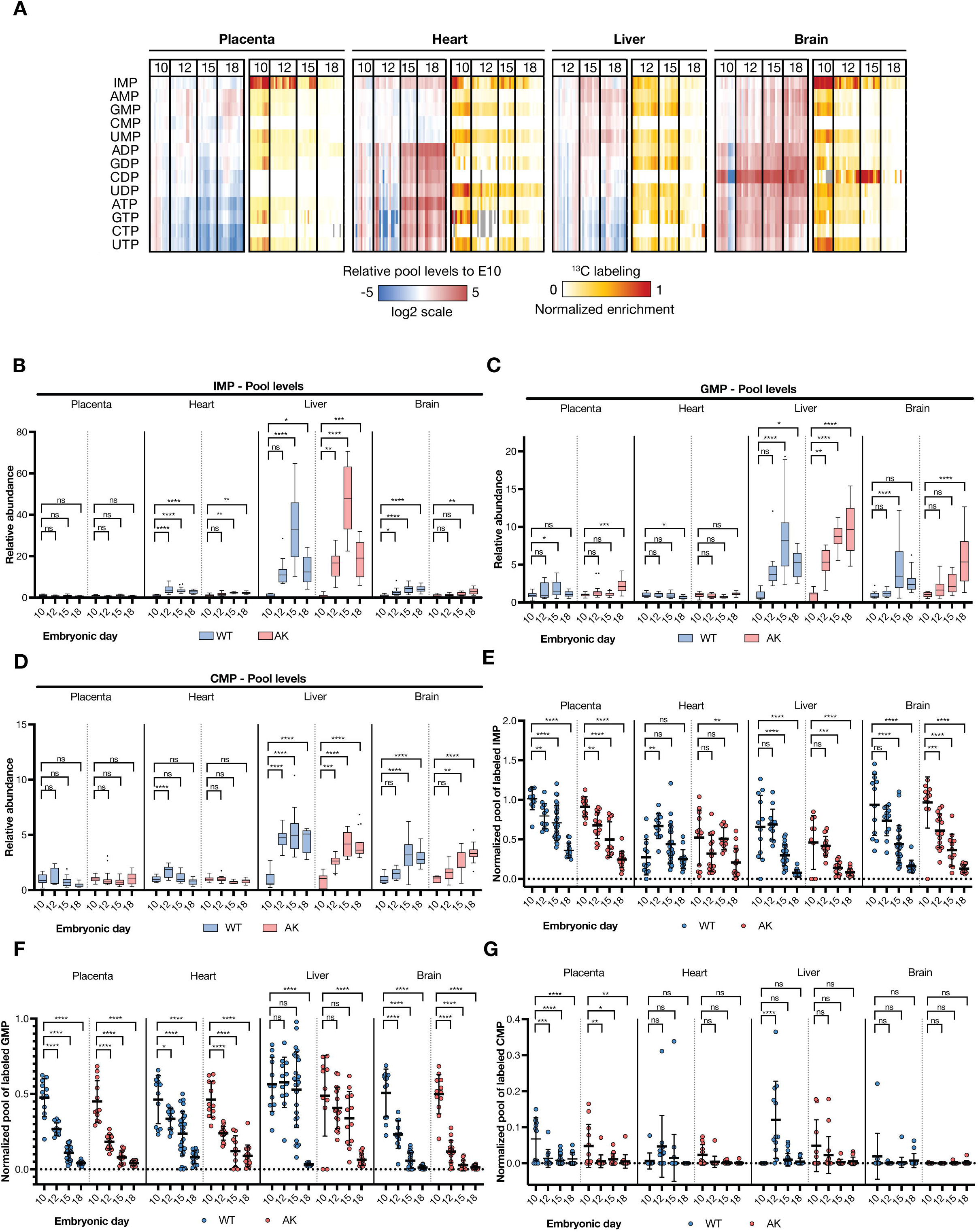
Progressive decrease in [U-^13^C_6_] glucose enrichment in fetal tissue nucleotides, related to Figure 5. (A) Blue to red scale: Heatmap representing the log2 fold changes in nucleotide pool levels relative to E10.5 fetal tissues from Akita dams. White to red scale: Heatmap representing the fractional pool of labeled nucleotides in fetal tissues from Akita during mid-to-late gestation. (B-D) Relative IMP (B), GMP (C), and CMP (D) pool levels across fetal tissues. For fetal tissues from WT dams, data is represented relative to the mean of WT E10.5 samples. For fetal tissues from Akita (AK) dams, data is represented relative to the mean of AK E10.5 samples. (E-G) Normalized pool of ^13^C-labeled IMP (E), GMP (F), and CMP (G) across fetal tissues. For each fetal tissue sample, labeling was normalized to the fractional pool of labeled glucose from its respective maternal labeling in plasma.

**Figure S5.**
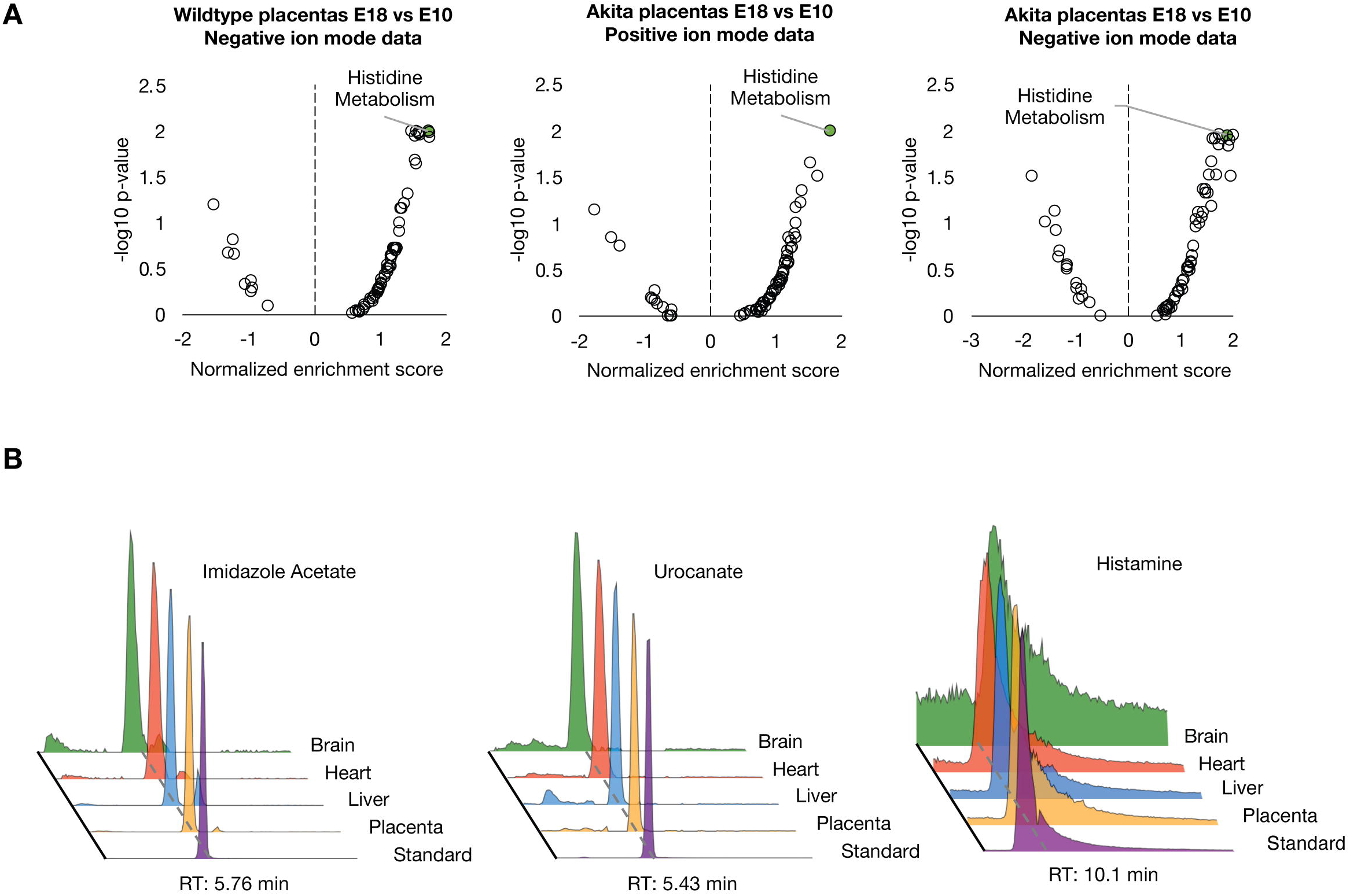
Untargeted analysis reveals increased levels of histidine derived metabolites in late gestation, related to Figure 6. (A) MetaboAnalyst functional analysis of negative ion mode untargeted data of E18.5 placentas from wildtype dams relative to E10.5 (left graph). MetaboAnalyst functional analysis of positive ion mode (middle graph) and negative ion mode (right graph) untargeted data of E18.5 placentas from Akita dams relative to E10.5. (B) Extracted ion chromatographs of imidazole-4-acetate, urocanate and histamine from representative fetal tissue samples compared to pure chemical standard. RT denotes retention time. Representative fetal tissue samples shown were from a E18.5 fetus (#1) of a wildtype dam (Mouse A).

## SUPPLEMENTAL TABLES

**Table S1. Targeted metabolomics of fetal tissues and maternal plasma, related to Figure 1**

**Table S2. Metabolite isotopologue distribution from fetal tissues and maternal plasma, related to Figure 1**

**Table S3. Untargeted metabolomics in placenta E18.5 and E10.5, related to Figure 6**

